# SARS-CoV-2 polyprotein substrate regulates the stepwise Mpro cleavage reaction

**DOI:** 10.1101/2022.09.09.507133

**Authors:** Manju Narwal, Jean-Paul Armache, Thomas Edwards, Katsuhiko S. Murakami

## Abstract

Processing of polyproteins pp1a and pp1ab in *Coronaviruses* by main protease Mpro is a crucial event in virus replication and a promising target for antiviral drug development. Mpro recognizes multiple recognition sites within polyproteins in a defined order, but its mechanism remains enigmatic due to lack of structural information of the polyprotein substrate bound Mpro complex. Here, we present the cryo-EM structures of the SARS-CoV-2 Mpro in an apo-form and in complex with the nsp7-10 region of pp1a polyprotein. The structure shows that the interaction of Mpro with polyproteins is limited to the recognition site connecting nsp9 and nsp10 proteins without any tight association with the rest of polyprotein structure or sequence. Comparison between the apo-form and the polyprotein bound structures of Mpro highlights the flexible nature of active site region allowing the Mpro to accommodate various recognition sites connecting series of nsp proteins. These observations suggest that the role of Mpro for selecting a preferred cleavage site within the polyprotein is limited and underscore the structure, conformation and/or dynamics of polyprotein determining the sequential polyprotein cleavage by Mpro.

## INTRODUCTION

Virus uses a wide variety of mechanisms for expressing genes by bypassing several requirements in protein translation in host cells such as having 5’ cap and poly-A tail on each mRNA. One of the most prevalent methods for expressing multiple proteins from a single mRNA frame is producing a precursor polyprotein followed by proteolytic cleavage (Hartenian et al., 2020). All the RNA and Retroviruses employ this strategy of translating their RNA genomes into long polyprotein chains. Polyprotein comprises of multiple proteins separated by protease recognition and digestion sites and produce multiple mature and functional proteins after cleavage at these sites. The polyprotein processing utilizes one or multiple viral and sometime host proteases in precise and highly regulated manner (Gupta, 2017). Since the polyprotein processing by viral protease is an essential step during virus replication in their hosts and the active sites of viral proteases are structurally distinct from their host proteases, the viral proteases such as HIV-1 protease in HIV and main protease in SARS-CoV-2 have been ideal targets for developing antiviral drugs against AIDS and COVID_19 treatments, respectively (Brik and Wong, 2003; Jin et al., 2020).

SARS-CoV-2 is a member of *Coronaviridae* family of viruses consisting of large positive sense single-stranded RNA genome of approximately 30kb in length with 5’ cap and 3’ poly-A tail for expressing non-structural proteins (nsps), and structural proteins are expressed by using subgenomic mRNAs (Hartenian *et al*., 2020). Two-third region of the virus RNA genome from 5’-end comprising of open reading frames ORF1 and ORF2 encoding pp1a and pp1b polyproteins, respectively. These polyproteins are subsequently processed into 16 individual nsps (nsp1-nsp16) to form replication/transcription complexes for the virus genomic RNA replication and the synthesis of subgenomic mRNAs. The polyproteins of SARS-CoV-2 are cleaved by the viral papain-like protease (PLpro) and the 3C-like cysteine protease (3CLpro) also known as Main protease (Mpro) (Gorkhali et al., 2021).

Mpro, encoded in nsp5, is responsible for its own release from the polyprotein through auto-proteolysis and then subsequently cleaves at 11 different sites found between nsp5 and nsp16 to release mature nsp protein. The Mpro recognition sites of SARS-CoV-2 consist of a consensus sequence of Q_(S/A/G/N) at P1_P1’ positions, which is highly conserved among different coronaviruses (Dai et al., 2020). Identification and cleavage of different recognition sequences by Mpro has been studied through the *in vitro* assays and X-ray crystallographic studies (Rut et al., 2020);. Crystal structures of the Mpro with recognition site sequence peptides highlighted the critical interactions between the Mpro substrate binding cleft and recognition site sequence, which explained substrate recognition and elucidated the mechanism of proteolysis (Lee et al., 2020) (MacDonald et al., 2021) (Zhao et al., 2022). Changing Mpro recognition sequence not only at P1_P1’ positions but also at flanking P2 or P2’ positions affect the efficiency of Mpro cleavage (Zhu et al., 2011) (Rut et al., 2021).

Functional nsps can be released from the polyproteins co-translationally. However, presence of different polyprotein intermediates during Murine Hepatitis Virus (MHV) infection indicates that polyproteins are processed by proteases post-translationally (Bost et al., 2000)) (Deming et al., 2007). The cleavage order of SARS-CoV polyprotein nsp7-10 region investigated by limited proteolysis and native mass spectrometry (native MS) revealed a stepwise cleavage; among three cleavage sites found in the nsp7-10 poly protein, the nsp9/10 site is cleaved first followed by cleavage at the subsequent sites (nsp7/8 and nsp8/9 sites) (Krichel et al., 2020). Amino acid sequences of the nsp7-10 region and Mpro are highly conserved (similarities are 98.5% and 96%, respectively), and the cleavage site sequences within the nsp7-10 region are identical in these species.

X-ray crystallographic studies of the SARS-CoV-2 Mpro has played a major role in understanding the mechanism of polyprotein cleavage and the structure-based drug design (Mengist et al., 2020). However, the structure of Mpro in complex with polyprotein remains undetermined due to flexible and unstable nature of polyprotein, which is not amenable for crystallographic study. In comparison to X-ray crystallography, cryo-electron microscopy (cryo-EM) can successfully investigate structures of flexible macromolecules (Xiao-chen Bai 2014).

In this study, we analyzed the order of nsp cleavage and release from polyprotein by Mpro through limited proteolysis assay and investigated the mechanism of stepwise polyprotein cleavage by determining the cryo-EM structures of the SARS-CoV-2 Mpro in the absence and presence of polyprotein substrate. We used the proteins containing nsp7-10 regions as a representative polyprotein instead of using its viral polyprotein substrates (pp1a and pp1a/b) since the nsp7-10 can be prepared homogeneously by bacterial expression system. Based on the structural findings and results of proteolysis assay, we propose that the polyprotein is the major determinant for sequential cleavage process by Mpro.

## RESULTS

### Polyprotein nsp7-10 processing by SARS-CoV-2 Mpro protease

To analyze the order of processing for the nsp7-10 region in SARS-CoV-2, we tested cleavage by Mpro through limited proteolysis. Addition of ZnCl_2_ in bacterial growth media during protein expression and maintaining high salt concentration during protein purification (Fig. 1A) facilitated the preparation of homogeneous nsp7-10 sample essential for the proteolysis assay and the structure study as described later. The reaction was initiated by mixing equal amount of nsp7-10 and Mpro (1:1 ratio), and all products including fully cleaved nsps and intermediates were analyzed on SDS-PAGE gel (Fig. 1B). Mpro attacks and cleaves the nsp9/10 site first thus nsp10 was released from the polyprotein (0.2 min), which simultaneously appeared with an intermediate nsp7-9 protein. As the reaction proceeded, nsp7-8 intermediate and nsp7, nsp8 and nsp9 products were released and the polyprotein processing was completed after 1 hour. We also monitored the reaction under polyprotein excess condition (Mpro:nsp7-10 = 1:4) to confirm that the nsp10 is cleaved first from the nsp7-10 polyprotein by Mpro at early stage of the reaction (15-60 sec, Fig. 1C). The result is consistent with the SARS_CoV_1 nsp7-10 polyprotein cleavage study (Krichel *et al*., 2020); among three cleavage sites available in the nsp7-10 polyprotein, Mpro cleaves the nsp9/10 site prior to other sites (Fig. 1D).

**Figure 1.**
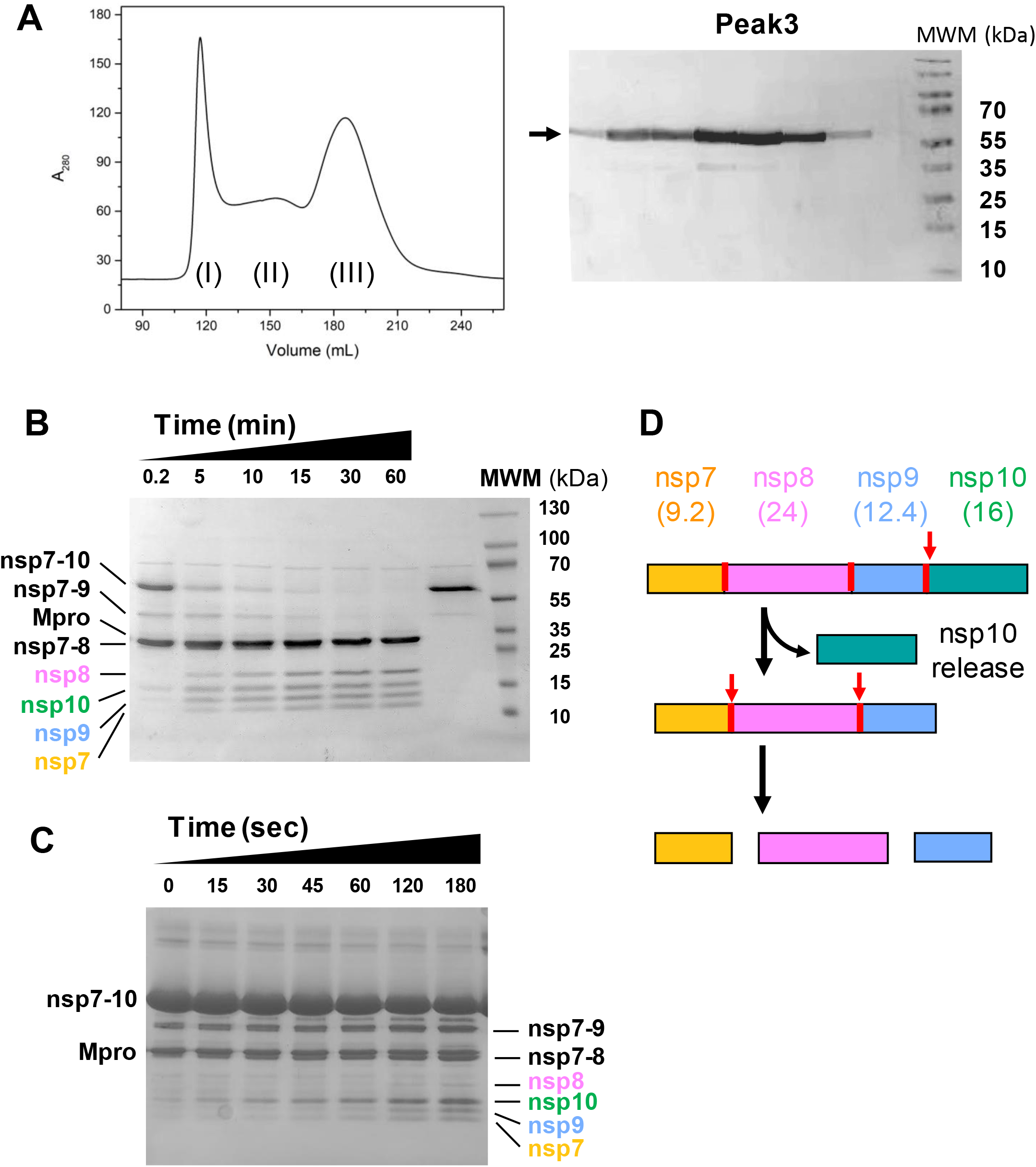
nsp7-10 polyprotein processing by Mpro. (A) Preparation of the nsp7-10 polyprotein. Different oligomers of nsp7-10 polyprotein (peaks I, II and III) are separated by size-exclusion chromatography in the presence of 1 M NaCl in the purification buffer (left). Proteins in fraction III are analyzed by SDS-PAGE (right). Molecular weight of the nsp7-10 polyprotein is 58.2 kDa. (B) Limited nsp7-10 polyprotein proteolysis by Mpro. Substrate (nsp7-10), products (nsp7, nsp8, nsp9 and nsp10), intermediates (nsp7-8, nsp7-9) and Mpro are separated by SDS-PAGE and labeled. Time after mixing Mpro with nsp7-10 (1:1 ratio) are indicated above lanes. (C) Limited nsp7-10 proteolysis assay under substrate excess condition (Mpro:nsp7-10 = 1:4). (D) Schematic illustration of the stepwise nsp7-10 polyprotein cleavage by Mpro. Non-structural proteins within the nsp7-10 and their molecular weights are indicated. Mpro recognition sites found between nsps are depicted as red lines and the cleavage order is indicated by red arrows.

### Cryo-EM structure of the Mpro with nsp7-10 polyprotein

To understand the structural basis of the cleavage site selection and the stepwise polyprotein processing by Mpro, we determined the 3D structure of Mpro and nsp7-10 polyprotein complex by cryo-EM single particle reconstruction. To form a stable complex of Mpro and nsp7-10, Cys145 residue of Mpro at its catalytic dyad residues was replaced to Ala (Mpro-C145A, Fig. S1A), which eliminates the protein cleavage without influencing its substrate binding at the Mpro active site (Ferreira et al., 2021; Narayanan et al., 2022).

Mpro-C145A was mixed with nsp7-10 at a molar ratio of 1:2 followed by isolating the complex through size-exclusion chromatography (Fig. S1B). To maintain the stability of the complex during sample vitrification, UltraAuFoil R1.2/1.3 were used, which also contributed to the high quality cryo-EM data collection by minimizing beam induced particle motions (Passmore and Russo, 2016). Cryo-EM micrographs were recorded at Titan Krios 300 kV microscope equipped with Gatan K3 summit direct electron detector and the structure was determined with an overall resolution of 2.49 Å (Figs. S2 and S3).

### Mpro contacts exclusively to the recognition site within the polyprotein

The cryo-EM map of the Mpro-C145A and nsp7-10 complex showed high-resolution features of the main and side chains of Mpro (domain I, residues 8-101; domain II, residues 102-184; domain III, residues 201-303) (Figs. 2A and B). In sharp contrast, observation of the nsp7-10 polyprotein density map is limited to the Mpro recognition site (10 residues, from P6 to P4’ positions), which is accommodated in the substrate binding cleft of both protomers of Mpro dimer (Figs. 2A and C). The Mpro exclusively binds the polyprotein through the recognition site; from P1 to P6 residues on one side of the scissile bond and from P1’ to P4’ residue on the other (Fig. 2C). Among the three recognition sites in the nsp7-10 polyprotein (nsp7/8, nsp8/9 and nsp9/10 sites), we assigned the density found at the Mpro active site as the nsp9/10 site sequence (amino acid sequence ATVRLQ↓AGNA, ↓ indicates a scissile bond), based on the best model fitting to the density map in comparison to the two other possible sites (nsp7/8, NRATL↓QAIAS; nsp8/9, SAVKL↓QNNEL) (Fig. S4A), and result of the polyprotein processing assay (Fig. 1B). Density for the rest of the nsp7-10 polyprotein could not be traced, however, upon Gaussian-filtering to the cryo-EM density map, fragmented densities likely corresponding to the main body of polyprotein becomes visible above the active site of Mpro (Fig. 2D). The individually determined structures of nsp7, nsp8, nsp9 and nsp10 (Konkolova et al., 2020) (Wang et al., 2020) (Littler et al., 2020) (Lin et al., 2020) cannot be fitted to the density due to lack of structural detail recognized in such lower resolution and fragmented map. The poorly defined density of the main body of polyprotein in the complex with Mpro suggests that Mpro contact with polyprotein only through the recognition site.

**Figure 2.**
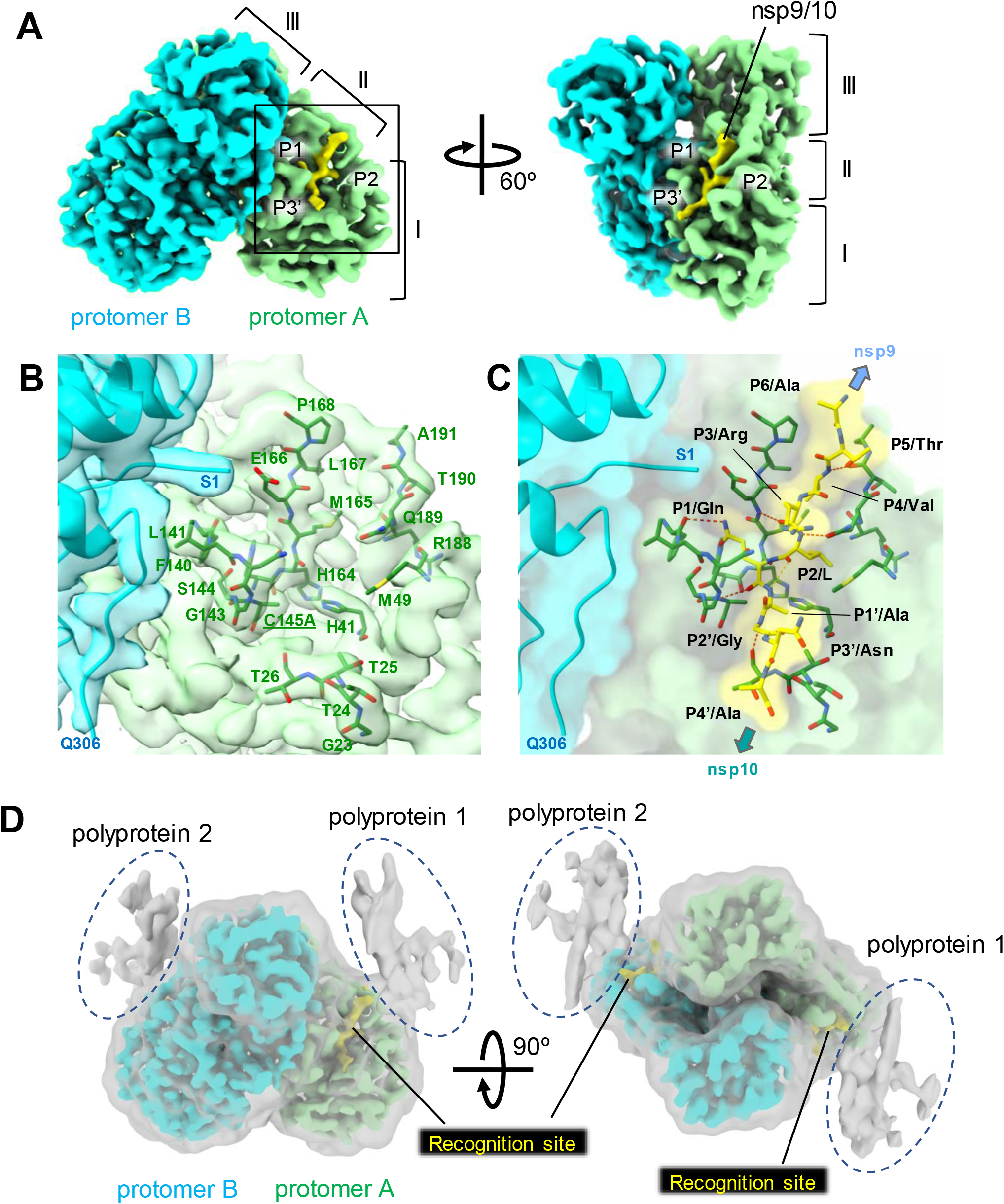
Cryo-EM structure of the nsp7-10 bound Mpro-C145A. (A) Cryo-EM density map of the Mpro-C145A and nsp7-10 complex. Density maps corresponding to each protomer of Mpro dimer and nsp9/10 recognition site are colored and indicated. Three domains of Mpro and P1, P2 and P3’ positions of the nsp9/10 are indicated. (B) A magnified view of the active center cleft of the Mpro (boxed area of protomer A in panel A). Cryo-EM density map is partially transparent and amino acid residues contacting to the nsp9/10 region (< 4 Å) are depicted as stick models and labeled. Density map and the model of nsp9/10 are removed to clarify. (C) A magnified view of the nsp9/10 and Mpro interaction. Structures of the nsp9/10 and Mpro residues participating in the nsp9/10 interaction are depicted as stick models with partially transparent surface model of the Mpro and nsp7-10 complex. Locations of the nsp9 and nsp10 are indicated. Orientation is the same as B. (D) Low path filtered cryo-EM density map of the Mpro and nsp9/10 complex (gray transparent overlayed with Mpro and nsp9/10 recognition site) shows the densities of the nsp7-10 polyprotein outside from the nsp9/10 region (dashed ovals). These densities locate above the active center of Mpro without any contact with Mpro.

The nsp9/10 site is accommodated in the active site cavity of Mpro extending along the substrate binding cleft forming antiparallel β strands (Fig. 2C). Mpro interacts with the P4 to P4’ position residues of the recognition site of polyprotein substrate by forming hydrogen bonds with the main chain of substrate. In addition, side chains of the substrate at P4/Val, P2/Leu, P1/Gln, P1’/Ala and P3’/Asn positions form the hydrogen bonds and/or van der Waals (vdW) interactions with Mpro. The three residues (P4, P2 and P’1) of the N-terminal (or non-prime) side of the cleavage site sequence fits into the narrow cavity of Mpro catalytic center. The glutamine at P1 (P1/Gln) rotated towards the oxyanion hole residues (Gly143 and Ser144) is positioned in the manner usually accepted to stabilize the scissile bond residues in Mpro and the P2/Leu is surrounded by H41, M49, H164, M165 and Q189 residues. The side chain of P3/Arg is surface exposed while carbonyl oxygen interaction with the Glu166 helps in its positioning close to the S1 site. The P4 to P6 residues of the recognition site are positioned near the S4 site, where main chain at P4 is making H-bonding contacts with Thr190. In contrast, the C-terminal (prime) side of the sequence is surface exposed except P1’ which is buried near the S1’ subsite. The Asn residue at P3’ orient to align and interact with T24 and Thr25 residues.

Binding of the nsp9/10 site at the Mpro active center is similar to the X-ray structures of the Mpro-C145A in complex with peptides containing nsp8/9 site (PDB:7MGR) and nsp4/5 site (PDB:7MGS) (Fig. S4B) (MacDonald *et al*., 2021). The similarity between the polyprotein substrate and the peptide substrate at the Mpro active center underscores that the Mpro contacts exclusively to the recognition site within the polyprotein.

### Cryo-EM structure of the apo-form Mpro

In addition to the Mpro-C145A and nsp7-10 polyprotein complex structure, we determined the cryo-EM structure of the wild-type Mpro in absence of substrate with an overall resolution of 3.7 Å (Figs. 3A and S5). We vitrified the Mpro on UltraAuFoil 1.2/1.3 grids, collected cryo-EM images and processed data as the Mpro-C145A and polyprotein complex (supplementary and material methods). The cryo-EM structure of Mpro is similar to the cryo-EM structure of Mpro-C145A and polyprotein complex determined in this study (RMSD: 0.604 Å), with noticeable differences at the active center near the P2 loop (residues T45 to L57) and P5 loop (residues from V186 to Q192) (Fig. 3B). Although the substrate binding at the Mpro active center does not trigger any major conformational change of Mpro, it reduces flexibility of the active center as indicated by moving P2 and P5 loops toward the active center and reducing B-factor of the P5 loop and domain I including the P2 loop in the Mpro and nsp7-10 complex compare with the apoform Mpro (Fig. 3C). B-factor of domain III is also reduced in the Mpro and nsp7-10 complex although the domain III does not participate in forming the substrate binding cavity of Mpro. Flexible nature of the active center and domain III of Mpro contribute a unique function of Mpro active center, which binds 13 different substrates in the pp1a and pp1ab polyproteins for processing.

**Figure 3.**
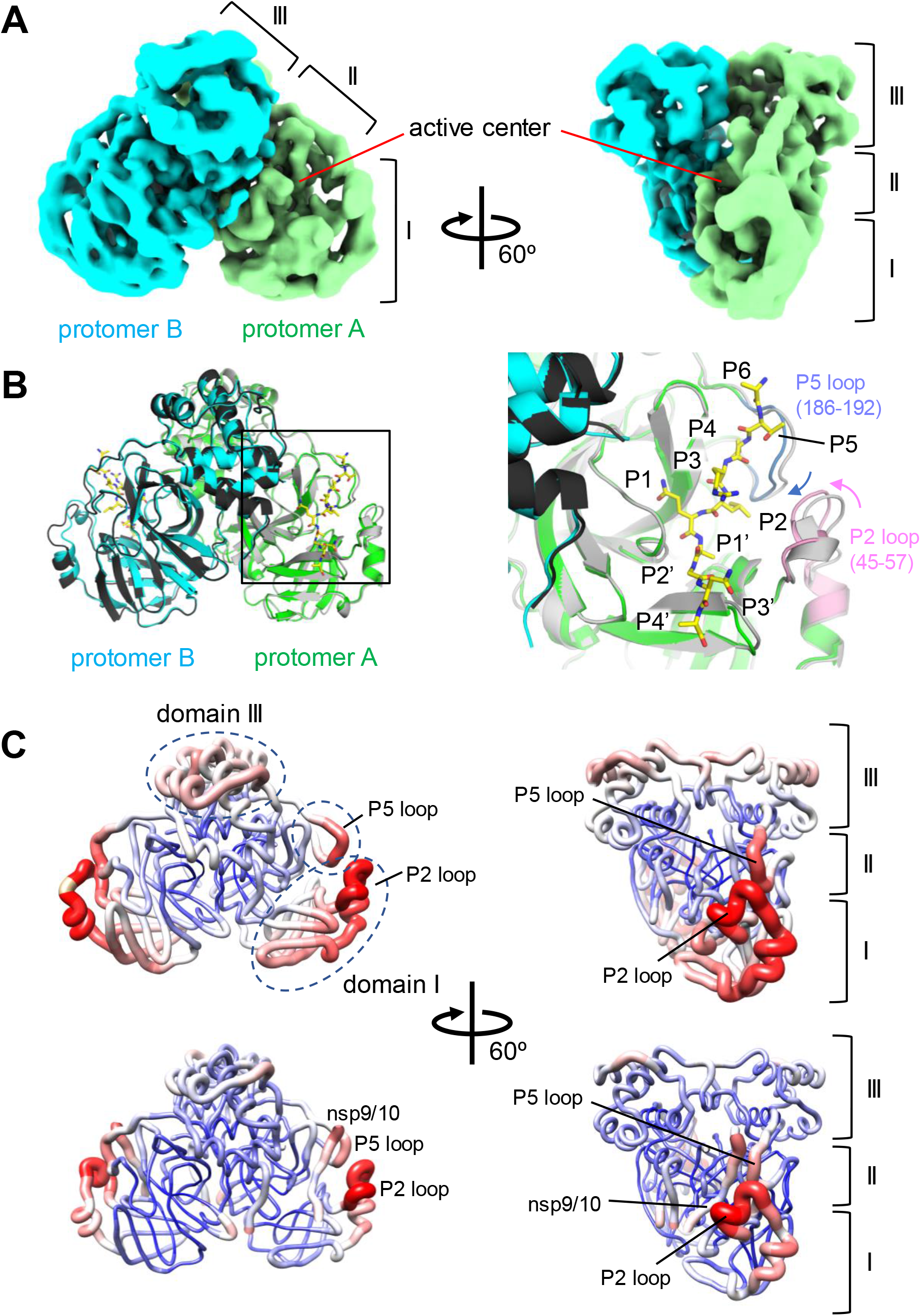
Cryo-EM structure of the wild-type Mpro. (A) Cryo-EM density map of the Mpro. Density maps corresponding to each protomer of Mpro dimer are colored and indicated. Three domains and active center of Mpro are indicated. (B) (left) Comparison of the Mpro structures in the apo-form (gray and block) and in the nsp7-10 complex (green, cyan and yellow). (right) A magnified view of the active center of the Mpro (boxed area of protomer A in right panel). The flexible P2 (pink) and P5 loops (blue) around the substrate binding cleft move toward the nsp9/10 recognition site in the Mpro and polyprotein complex (indicated by pink and blue arrows). (C) Comparison of the B-factor distributions in the apo-form (top) and in the nsp7-10 complex (bottom) of Mpro. Cartoon representation of the models with gradient of color (blue, white to red) and thickness (narrow to wide) reflecting the scale of the B factors (low to high). Domains and loops of the apo-form Mpro showing higher B-factor compare with the Mpro and nsp7/10 complex are indicated by black dashed ovals.

## DISCUSSION

Since the complete genome sequence of SARS-CoV-2 was posted online on early January 2020, the structure of Mpro and substrate complex has been extensively investigated using recognition peptide as a substrate for understanding the mechanism of enzymatic action. However, the interaction between Mpro and polyprotein and the mechanism of polyprotein processing have not yet been investigated in detail. This study shed light on the Mpro and polyprotein interaction and the mode of cleavage and release of nsp proteins from polyprotein.

We used the SARS-CoV-2 polyprotein containing the nsp7-10 region as a model substrate since this region is highly conserved among *Coronaviruses* and these nsp proteins are critical for effective viral replication in infected cells. The Mpro cleaves the nsp7-10 polyprotein stepwise, starting at the nsp9/10 site first followed by cleaving the nsp8/9 and nsp7/8 sites (Figs. 1B, C and D). We also determined the cryo-EM structures of the Mpro in the presence and in the absence of the polyprotein substrate for the first time (Figs. 2 and 3), which revealed that Mpro strongly associates with the recognition site residues but does not contact with the rest of polyprotein structure or sequences (Fig. 2D).

After releasing itself from the polyprotein by autocleavage reaction, the Mpro cleaves polyproteins beginning from the C-terminus region at nsp9 (nsp9/10 site) in case of polyprotein region containing nsp7-10 although two additional cleavage sites at nsp7/8 and nsp8/9 found in this region. We propose two possible models explaining the stepwise polyprotein processing. In the first “polyprotein driven model” (Fig. 4A), limited number of the recognition sites are exposed on surface of polyprotein for recruiting Mpro. Cleaving preferred recognition sites and releasing nsps exposes additional recognition sites that recruit Mpro to continue processing. In this model, cleavage order is determined by polyprotein and Mpro plays a limited role in determining the order of cleavage reaction. In the second “Mpro driven model” (Fig. 4B), most recognition sites are exposed on surface of the polyprotein and Mpro selects a preferred cleavage site based on its affinity to the Mpro. Since recognition sequences are highly conserved, in this model, Mpro may contact the polyprotein beyond the recognition site to make energetically favored associations of Mpro with one of the cleavage sites over others. The results from this study support the polyprotein driven model (Fig. 4A) since the cryo-EM study of the Mpro and polyprotein complex showed that the Mpro forms a complex with polyprotein only through the recognition site. Complete understanding the mechanism of stepwise processing of polyprotein requires further structural and biochemical investigation of the Mpro and polyprotein interaction and processing.

**Figure 4.**
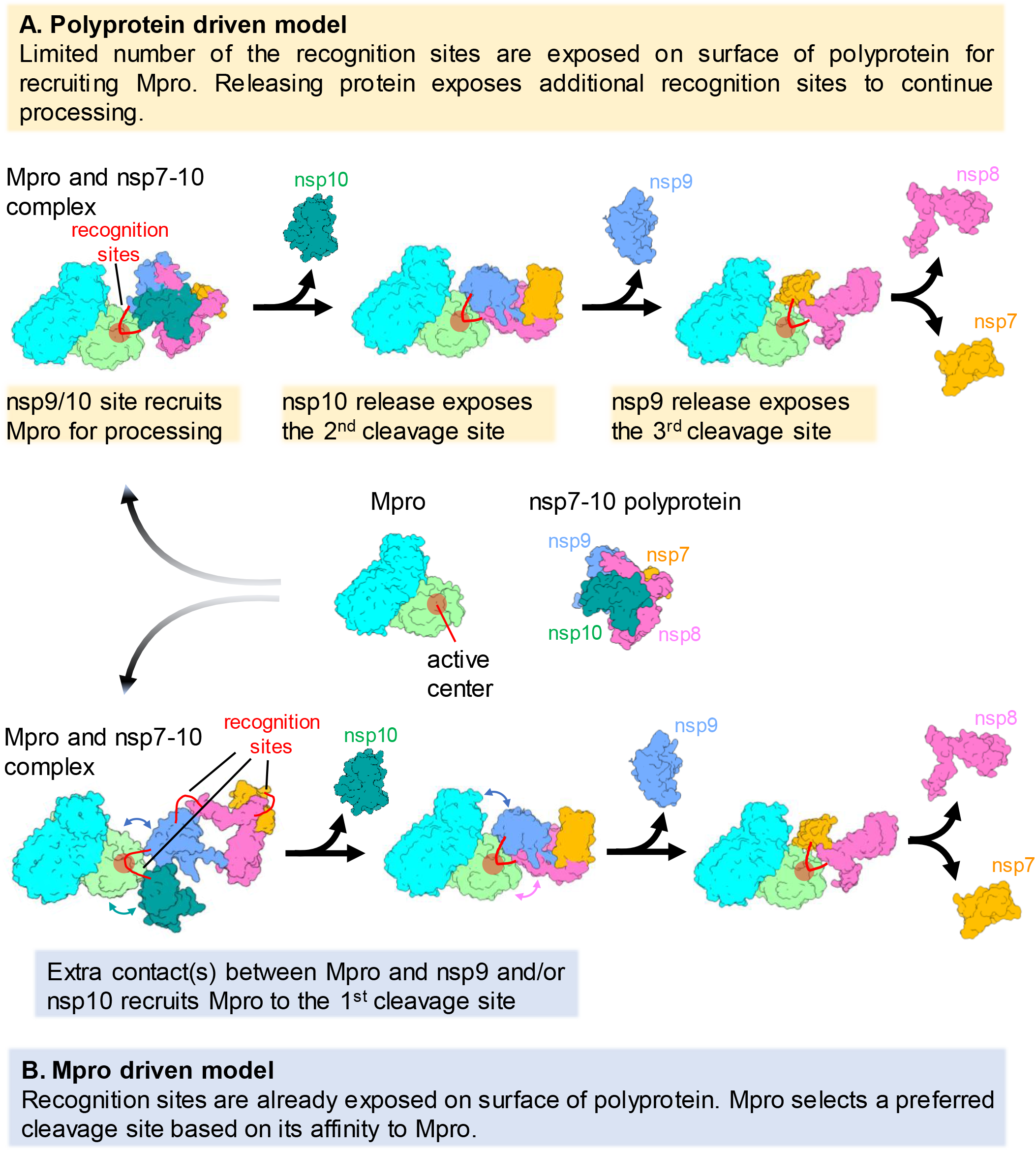
Two distinct models explaining the stepwise polyprotein process by Mpro.

Although “resolution revolution” of cryo-EM in the last decade, determining atomic-resolution structure of small macromolecules (less than 100 kDa molecular weight) by cryo-EM single-particle reconstruction remains challenge as reflected in the fact that less than 20 such macromolecule structures determined by cryo-EM single particle reconstruction (with their resolutions higher than 3.0 Å) found in the Protein Data Base (PDB). By careful selection of grids for minimizing beam induced particle motion and extensive corrections and classifications during data processing made it possible for us to detremine the structure of Mpro at 2.5 Å resolution.

Conservation of the domain architecture in the cryo-EM structure of Mpro in comparison with the X-ray crystal structures of the Mpro confirms that the cryo-EM can be an alternative method for investigating the structure of Mpro. The cryo-EM study can provide additional valuable insight regarding the dynamics of substrate binding region of the Mpro (Fig. 3C). The X-ray crystallography cannot avoid some artifacts such as influencing structure and/or B-factor distribution of macromolecules due to packing interactions among symmetry related macromolecules (packing artifact) (Fig. S7A and B). In our previously determined X-ray crystal structure of the apo-form Mpro (PDB: 7LKD, 2.01 Å resolution) (Narayanan *et al*., 2022), B-factors around the active center of Mpro are not identical in each protomer (protomers A and B, Fig. S7B) due to their environments within the crystal (Fig. S7A).

In summary, we provide insights into how SARS-CoV-2 Mpro would recognize and cleave on multiple sites on polyprotein substrate. This study establishes the role of polyprotein towards the sequential order of processing by Mpro and lay the foundation for futures studies aimed at understanding the interactions of these proteins during the coronavirus replication.

## EXPERIMENTAL PROCEDURES

### Cloning, Protein Expression and Purification

Plasmid pSUMO is used for producing WT and derivative of SARS-Co-V2 Mpro as described (Narayanan *et al*., 2022). Recombinant Mpro was expressed in *E. coli* BL21(DE3) cells and purified by Ni-affinity and size-exclusion chromatography (Fig. S1A). The final purified protein in buffer 20 mM HEPES (pH 7.5), 5% glycerol, 100 mM NaCl and 0.5 mM DTT was stored at −80 °C for further use.

The codon-optimized gene encoding the nsp7-10 region of SARS-CoV-2 polyprotein was synthesized commercially (GenScript) and cloned between the *NdeI* and *XhoI* sites of pET15b vector, which introduces a N-terminal 6x His-tag. The nsp7-10 was expressed in BL21(DE3) cells in LB growth media containing 0.1 mM ZnCl_2_ along with ampicillin. Next day, 1000 mL media was inoculated with the 5 mL overnight grown culture. The protein overexpression was induced with 0.4 mM IPTG at OD_600_ 0.6-0.7 and grown for 16h at 16 °C.

The cells were pelleted by centrifugation at 7000 g for 20 min (4 °C) and resuspended in 100 ml of lysis buffer (20 mM Tris-HCl pH 8.0, 5% glycerol, 500 mM NaCl and 1 mM DTT). Cells were disrupted by sonication (60% amplitude, for 5min; pulse 5s on/10s off), after which the lysate was cleared by centrifugation (39,000 g, 30 min, 4 °C).

The polyprotein was purified after loading the supernatant on a HisTrap HP column equllibrated with buffer A (20 mM Tris pH 8.0, 500mM NaCl, 5% Glycerol, 1mM DTT) and eluting the protein after washing with a gadient with buffer B ((20 mM Tris pH 8.0, 200mM NaCl, 5% Glycerol, 300mM Imidazole, 1mM DTT). Protein-containing fractions were diluted 4 times with buffer C (20 mM Tris pH 8.0, 5% Glycerol, 1mM DTT) and loaded on Hitrap Q column. The protein was eluted with 50% buffer D (20mM tris pH 8.0, 1M NaCl, 5% Glycerol, 1mM DTT) and loaded on Superdex 200 26/600 pre-equilibrated with buffer E (20mM HEPES pH 7.5, 10% Glycerol, 1M NaCl and 5mM DTT) (Fig. 1A).

### Polyprotein processing assay

The nsp7-10 polyprotein processing by the Mpro protease was monitored in an *in vitro* assay (Figs. 1B and 1C). The reaction was set-up in the assay buffer (20 mM HEPES pH 7.5, 5% glycerol, 1 mM DTT, 100 mM NaCl) by incubating wild-type Mpro and nsp7-10 polyprotein in a molar ratio of 1:1 (Fig. 1B) or 1:4 (Fig. 1C) at room temperature. A total reaction volume of 150 μL was prepared which included adding the polyprotein in the assay buffer followed by the addition of WT Mpro protease. From this reaction mixture, 10 μL aliquots were taken at regular time intervals starting from 0 to 60 min and mixed with Laemmli sample buffer to stop the reaction (4 volumes, 1× final). The aliquoted reaction mixtures were denatured (60 °C, 10 min) and analyzed by SDS-PAGE.

### Cryo-EM sample preparation, data collection and processing

To prepare the Mpro and nsp7-10 complex for the cryo-EM data collection, the Mpro-C145A (5 mg/ml) and nsp7-10 polyprotein (9 mg/ml) were mixed (1:2 molar ratio, 500 microL volume) and pre-incubated overnight at 4 °C in solution containing 20 mM HEPES (pH 7.5), 100 mM NaCl, 0.5 mM DTT, 5 % glycerol. The complex was loaded on a Superose 6 10/300 column (GE healthcare, currently Cytiva) pre-equilibrated with 20 mM HEPES (pH 7.5), 100 mM NaCl, 0.5 mM DTT buffer and isolated from a single peak (Fig. S1B). A 3.5 μL sample containing the complex (1 mg/ml) was applied to a glow-discharged UltraAuFoil grid (R1.2/1.3, mesh 300) (Electron Microscopy Sciences), and then blotted and plunge-frozen in liquid ethane using a Vitrobot Mark IV (FEI, USA) with 95 % humidity at 4 °C. The cryo-EM grids for the apo-form wild-type Mpro was prepared as described above. The cryo-EM grids were imaged using a 300 keV Titan Krios (Thermo Fisher) microscope equipped with a K3 direct electron detector (Gatan) and controlled by the Latitude S (Gatan, Inc.) software at the National Cancer Institute Cryo-EM Facility at Frederick. The defocus range for data collection was −1.0 to −2.5 μm, and the magnification was × 105,000 in electron counting mode (pixel size = 0.873 Å/pixel). Forty frames per movie were collected with a dose of 1.125 e^-^/Å^2^/frame, giving a total dose of 50 e-/Å^2^.

The MproC145A-nsp710 polyprotein complex data were processed using cryoSPARC (Punjani et al., 2017) (Figs. S2 and 3). A total of 5,606 movies were collected, and the movies were aligned, and dose weighted using Patch-motion correction. CTF fitting was performed with Patch-CTF estimation. Initially, ~1000 particles were manually picked to generate particle templates followed by automated picking, resulting in a total of 418,049 particles subjected to 2D classification. After two rounds of 2D classification to remove junk particles, 361,048 particles were used to generate two *ab initio* models. Junk particles were removed, resulting in a dataset of 275,629 particles chosen for the 3D classification (heterogenous refinement). Poorly populated classes were removed, resulting in a dataset of 49,995 particles to generate the density map at 2.49 Å resolution. The particles were 3D auto-refined without the mask and post-processed (homogenous refinement). The cryo-EM data of the apo-form wild-type Mpro was processed as the Mpro-C145A and nsp7-10 complex and outlined in Fig. S6.

### Model building and refinement

To refine the Mpro-C145A and nsp7-10 polyprotein structure, the X-ray crystal structure of Mpro (PDB: 7LBN) (Narayanan *et al*., 2022) was manually fit into the cryo-EM density map using Chimera (Pettersen et al., 2004). The structure of nsp7-10 recognition sequence was manually built, and real-space refined by using Coot (Emsley and Cowtan, 2004). The structure was real-space refined using Phenix (Afonine et al., 2010) with secondary structure, Ramachandran, rotamer, and reference model restraints. The structure of apo-form wild-type Mpro was refined as describe in the Mpro and nsp7-10 complex (Table S1). Figures were prepared by ChimeraX and Pymol.

## ACKNOWLEDGEMENTS

We thank Dr. Joyce Jose for providing a plasmid with Mpro encoding gene. We thank Carol Bator at Penn State for supporting initial screening and optimization of the cryo-EM specimen preparations. We thank Prof. Eddy Arnold at Rutgers University and Prof. Patrick Griffin at Scripps Research Institute, Florida for critical discussion of the manuscript. This research was, in part, supported by the National Cancer Institute’s National Cryo-EM Facility at the Frederick National Laboratory for Cancer Research under contract HSSN261200800001E. This work was supported by NIH grant (R35 GM131860 to K.S.M.).

---

**Figure S1.**
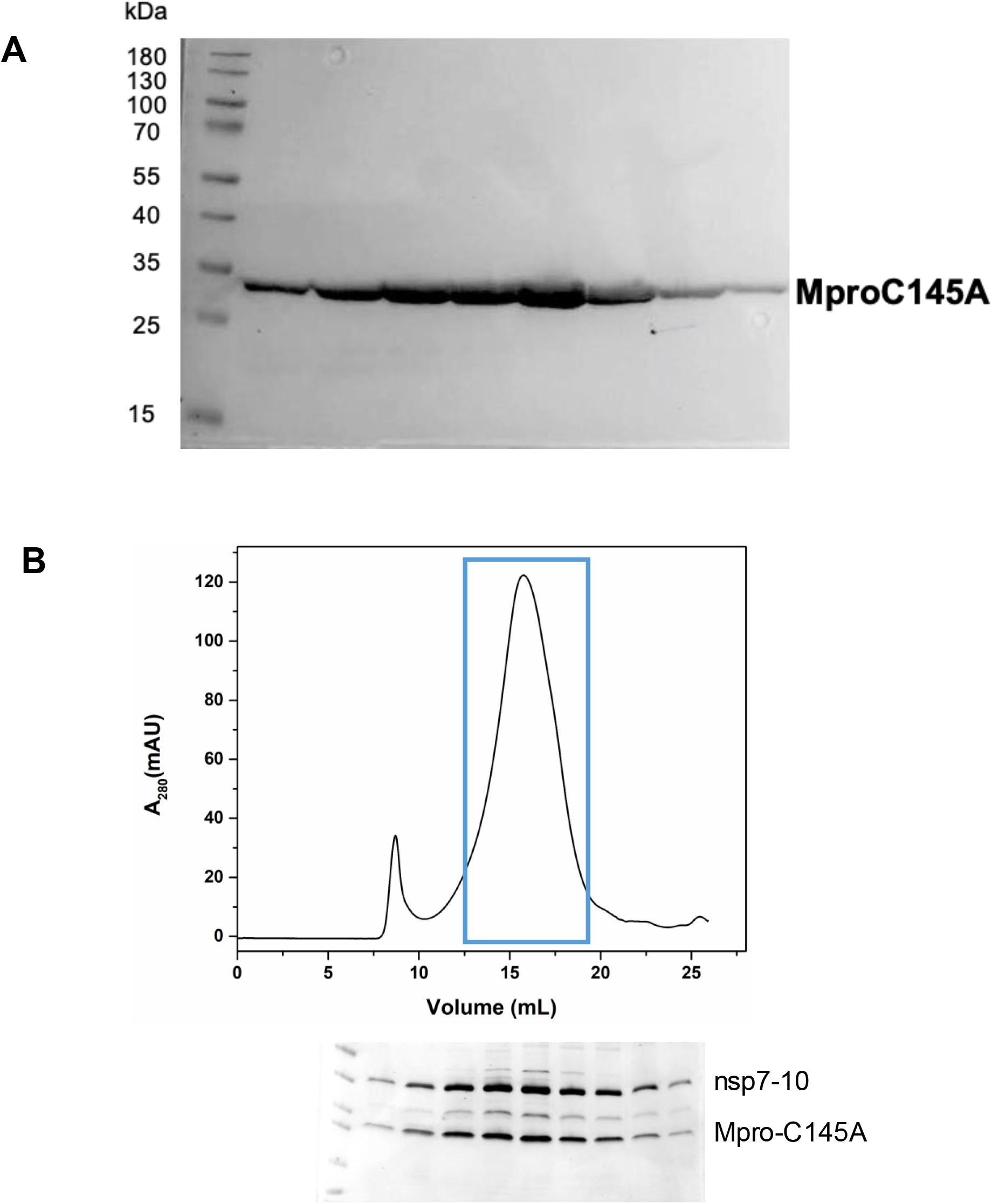
(A) Purification of Mpro-C145A by size exclusion chromatography. (B) Purification of the Mpro-C145Aand nsp7-10 complex by size exclusion chromatography (top). SDS-PAGE analysis confirmed peak fractions (blue line boxed) containing both Mpro and nsp7-10 (bottom).

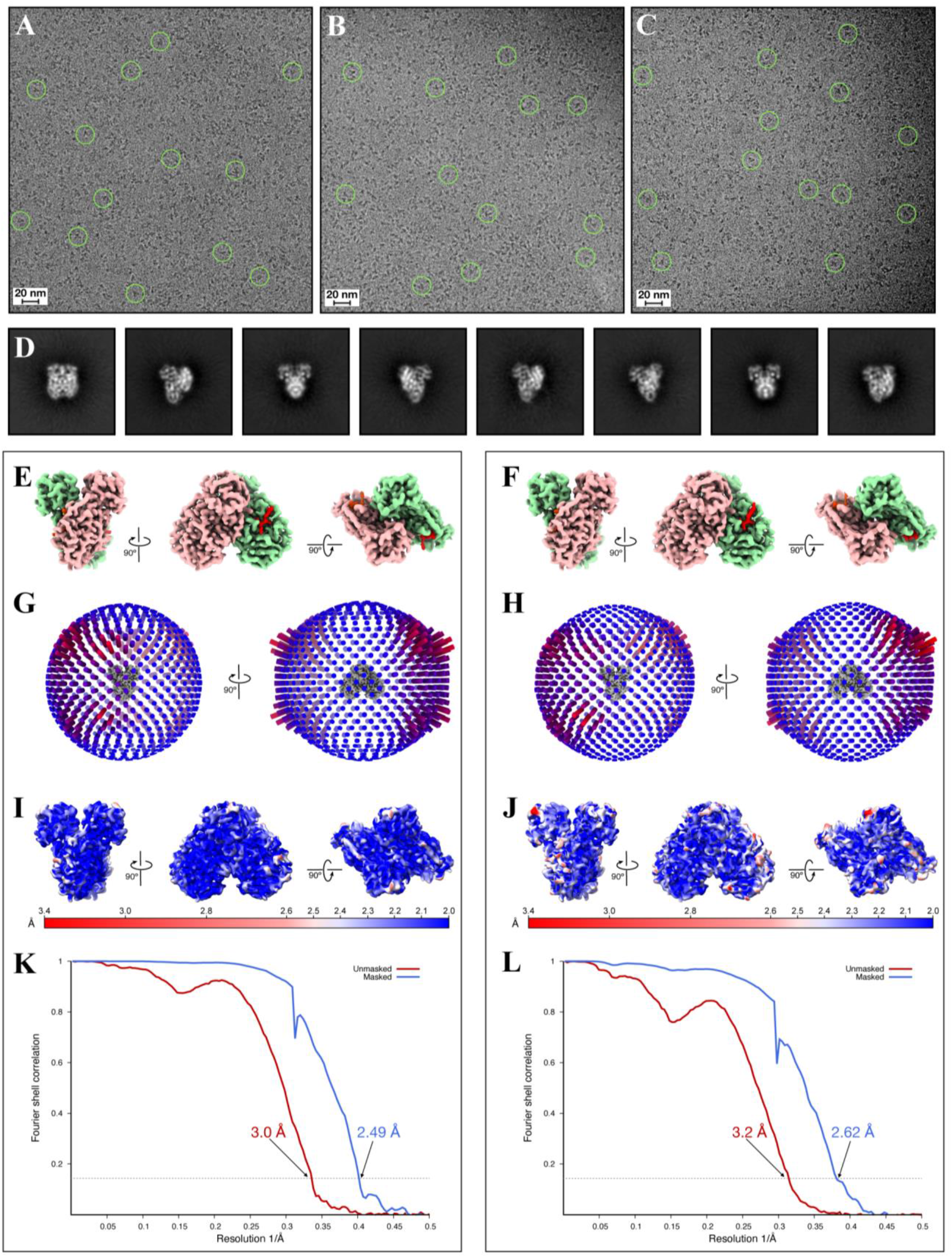

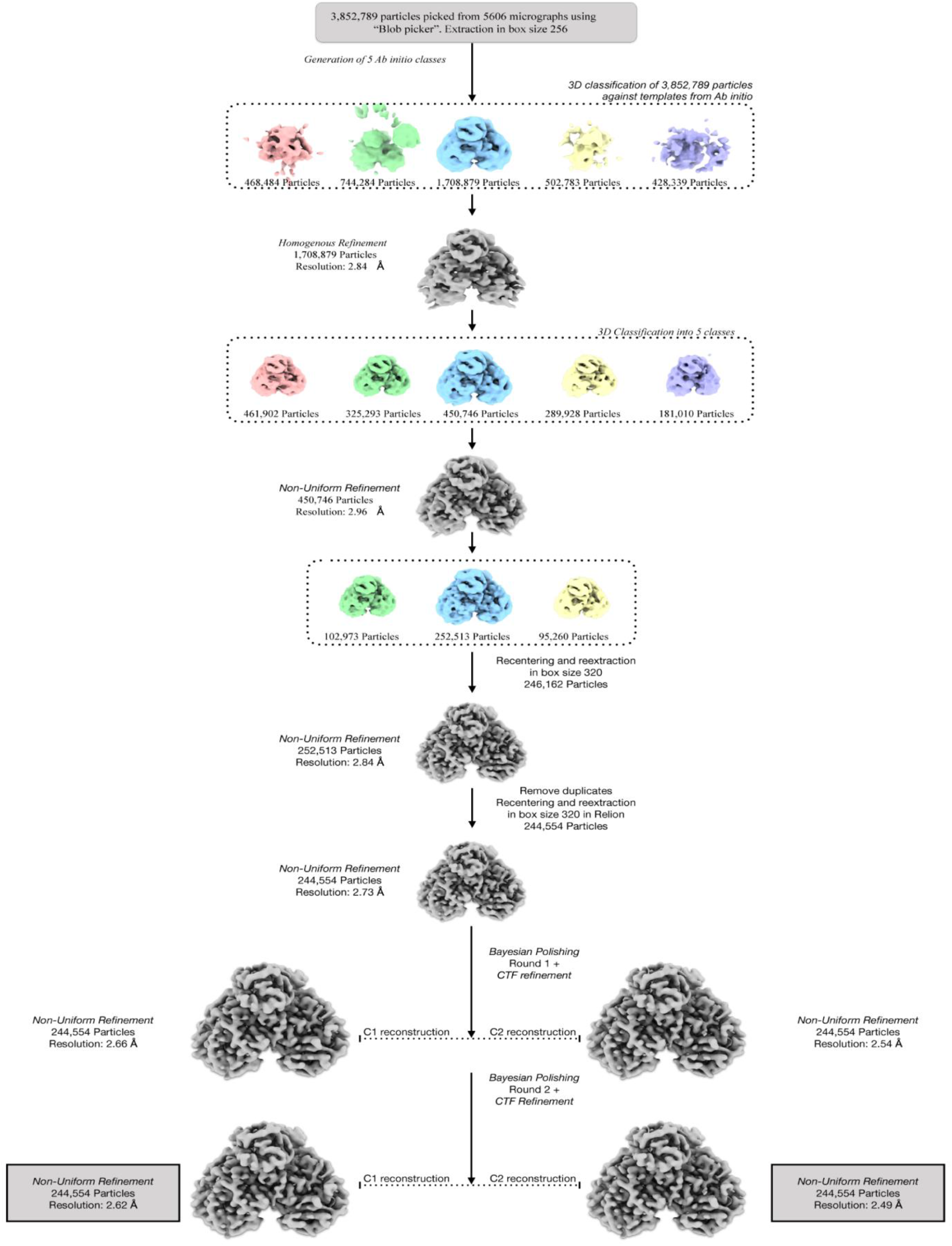

**Figure S4.**
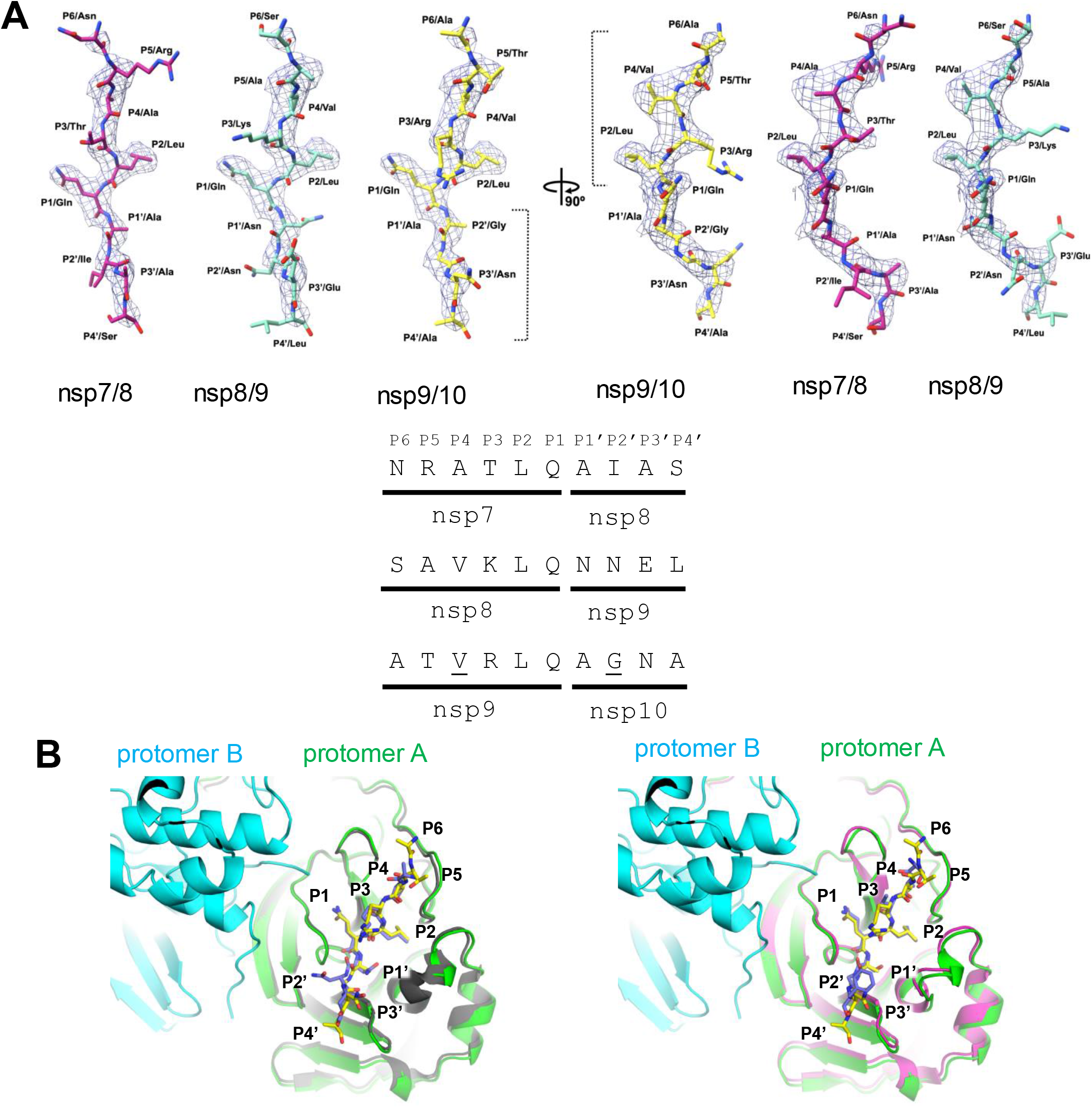
(A) Fitting the Mpro recognition sites, nsp7/8, nsp8/9 and nsp9/10, on the cryo-EM density map of the Mpro-C145A and nsp7-10 complex. At nsp8/9 and nsp7/8 sites, larger side chain amino acids i.e Asn and Ile respectively, present at P2’ positions are extending beyond the map density, whereas smaller Ala at P4 position in nsp7/8 rules out this as the possible site. Overall, variations in the residues at these positions makes nsp9/10 as the best fitting site in the map density among all the recognition sites on nsp7-10.(B) Substrate recognition site binding in the Mpro active site. Binding of nsp8/9 (colored light yellow) (PDB: 7MGR) and nsp4/5 (colored brown) (PDB:7MGS) in Mpro-C145A X-ray derived structure is similar to the nsp9/10 recognition site (colored yellow) in cryo-EM derived structure.

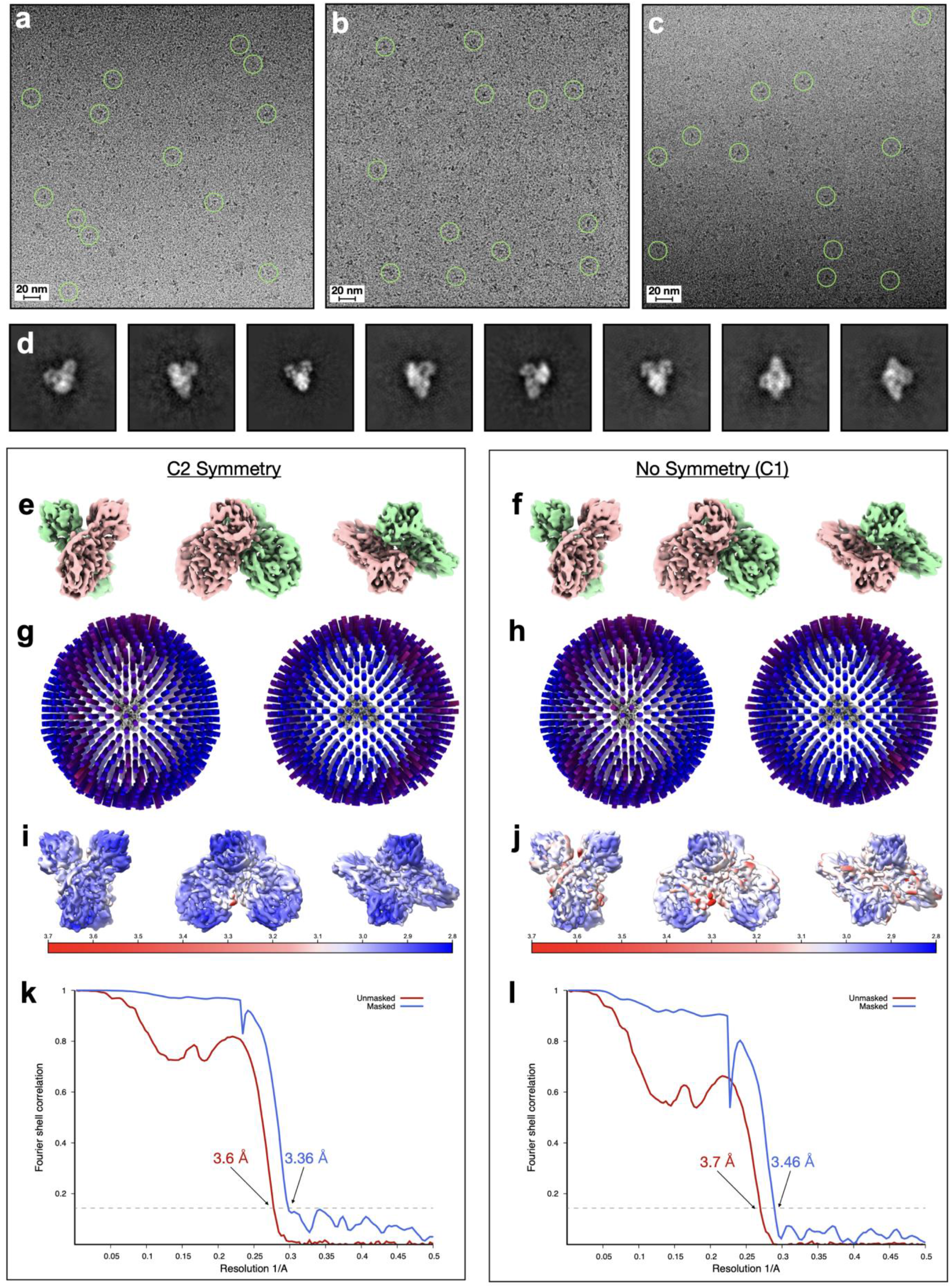

**Figure S6.**
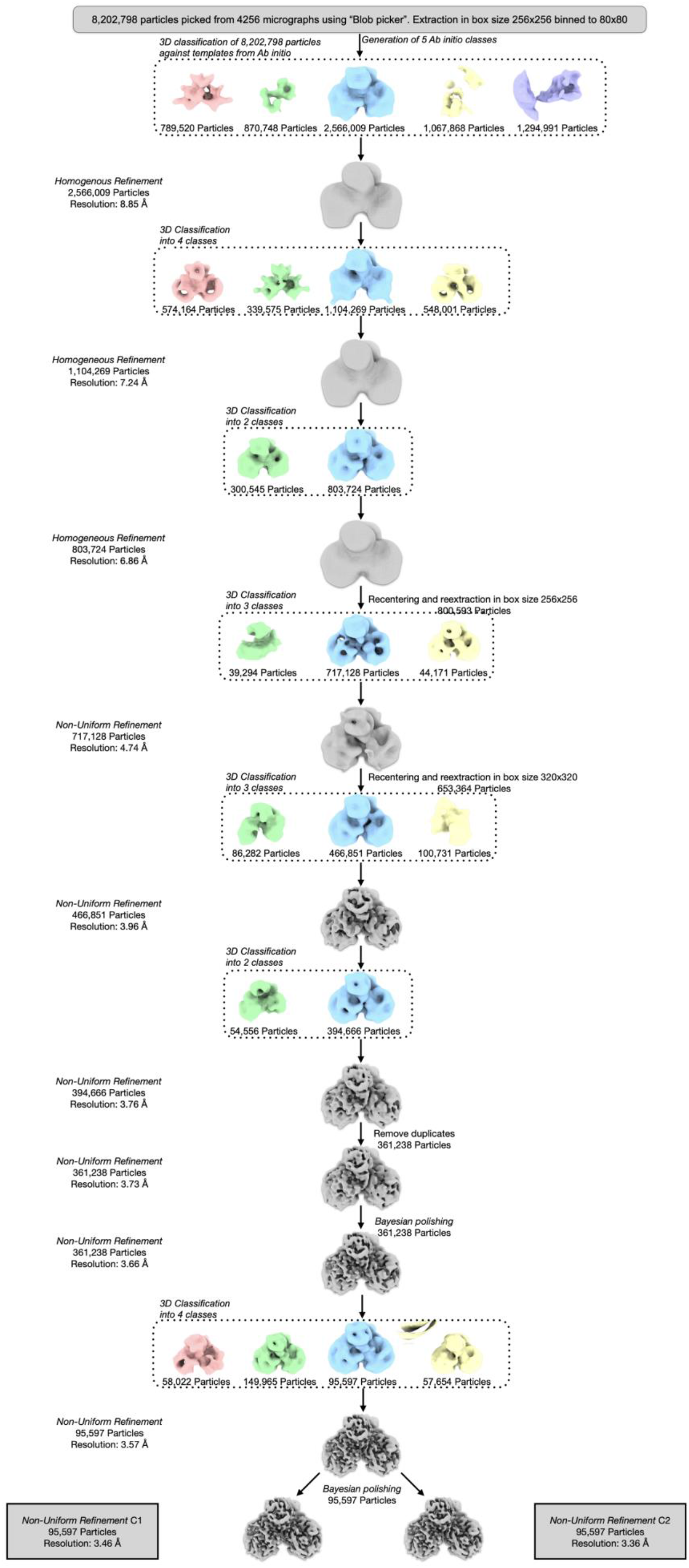
Crystal structure analysis of WT Mpro protease (PDB:7LKD (A) Threedimensional crystal packing of Mpro. The flexible P5 and P2 loop are limited due to the surrounding monomers packed in the crystal lattice. (B) B-factor values for the model of the WT protease are illustrated by color and thickness (red/thick for higher B-factor to blue/thin for lower). B-factor distribution in both the protomers in the model is uneven due to the packing effects of crystal, affecting the reliable interpretation of the model features for understanding their biological significance.

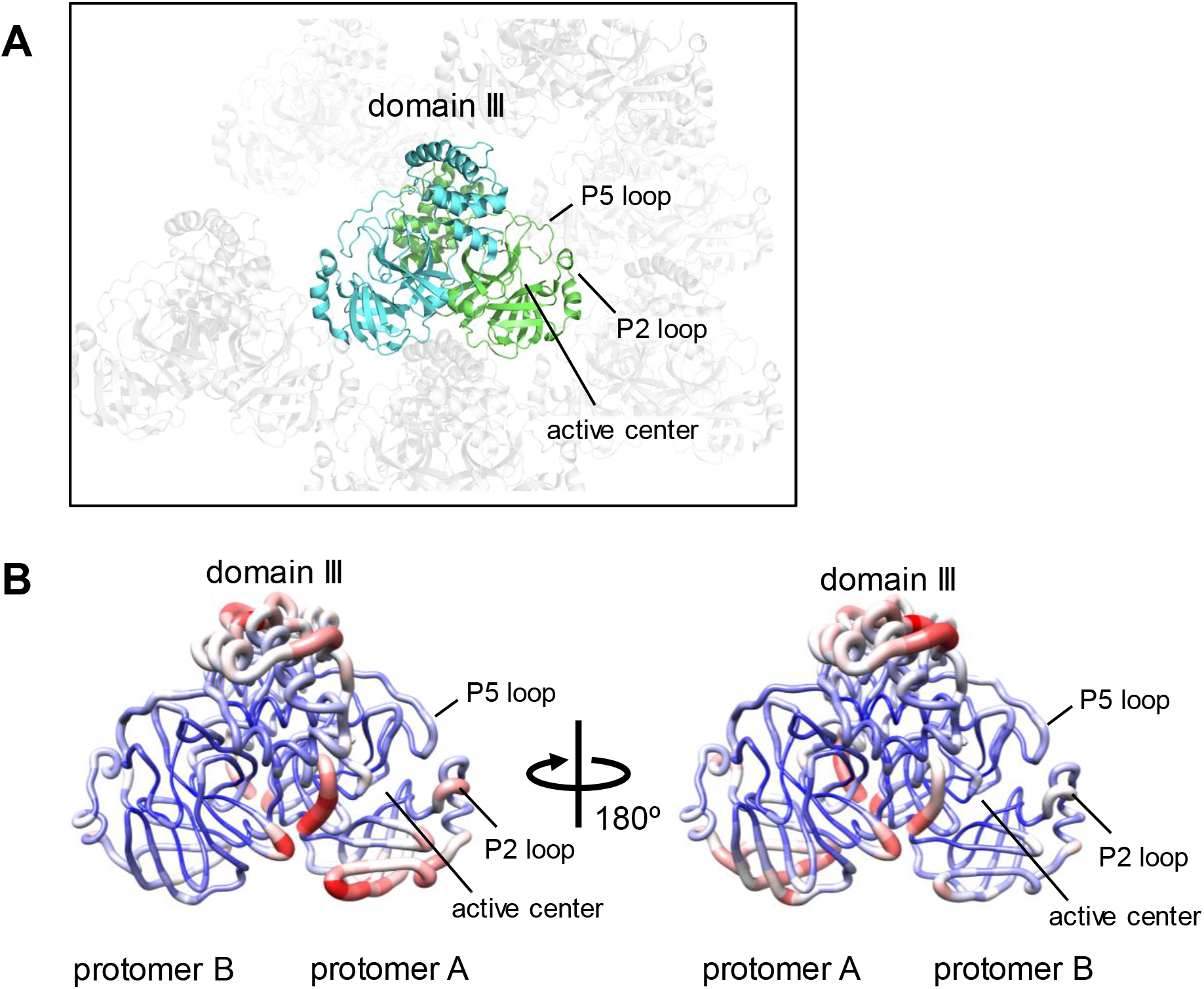

## REFERENCES

Afonine, P.V., Mustyakimov, M., Grosse-Kunstleve, R.W., Moriarty, N.W., Langan, P., and Adams, P.D. (2010). Joint X-ray and neutron refinement with phenix.refine. Acta crystallographica 66, 1153–1163. 10.1107/S0907444910026582.

Bost, A.G., Carnahan, R.H., Lu, X.T., and Denison, M.R. (2000). Four proteins processed from the replicase gene polyprotein of mouse hepatitis virus colocalize in the cell periphery and adjacent to sites of virion assembly. J Virol 74, 3379–3387. 10.1128/jvi.74.7.3379-3387.2000.

Brik, A., and Wong, C.H. (2003). HIV-1 protease: mechanism and drug discovery. Org Biomol Chem 1, 5–14. 10.1039/b208248a.

Dai, W., Zhang, B., Jiang, X.M., Su, H., Li, J., Zhao, Y., Xie, X., Jin, Z., Peng, J., Liu, F., et al. (2020). Structure-based design of antiviral drug candidates targeting the SARS-CoV-2 main protease. Science 368, 1331–1335. 10.1126/science.abb4489.

Deming, D.J., Graham, R.L., Denison, M.R., and Baric, R.S. (2007). Processing of open reading frame 1a replicase proteins nsp7 to nsp10 in murine hepatitis virus strain A59 replication. J Virol 81, 10280–10291. 10.1128/JVI.00017-07.

Emsley, P., and Cowtan, K. (2004). Coot: model-building tools for molecular graphics. Acta crystallographica 60, 2126–2132.

Ferreira, J.C., Fadl, S., Villanueva, A.J., and Rabeh, W.M. (2021). Catalytic Dyad Residues His41 and Cys145 Impact the Catalytic Activity and Overall Conformational Fold of the Main SARS-CoV-2 Protease 3-Chymotrypsin-Like Protease. Front Chem 9, 692168. 10.3389/fchem.2021.692168.

Gorkhali, R., Koirala, P., Rijal, S., Mainali, A., Baral, A., and Bhattarai, H.K. (2021). Structure and Function of Major SARS-CoV-2 and SARS-CoV Proteins. Bioinform Biol Insights 15, 11779322211025876. 10.1177/11779322211025876.

Gupta, S.P. (2017). Viral proteases and their inhibitors (Elsevier/Academic Press).

Hartenian, E., Nandakumar, D., Lari, A., Ly, M., Tucker, J.M., and Glaunsinger, B.A. (2020). The molecular virology of coronaviruses. J Biol Chem 295, 12910–12934. 10.1074/jbc.REV120.013930.

Jin, Z., Du, X., Xu, Y., Deng, Y., Liu, M., Zhao, Y., Zhang, B., Li, X., Zhang, L., Peng, C., et al. (2020). Structure of M(pro) from SARS-CoV-2 and discovery of its inhibitors. Nature 582, 289–293. 10.1038/s41586-020-2223-y.

Konkolova, E., Klima, M., Nencka, R., and Boura, E. (2020). Structural analysis of the putative SARS-CoV-2 primase complex. J Struct Biol 211, 107548. 10.1016/j.jsb.2020.107548.

Krichel, B., Falke, S., Hilgenfeld, R., Redecke, L., and Uetrecht, C. (2020). Processing of the SARS-CoV pp1a/ab nsp7-10 region. Biochem J 477, 1009–1019. 10.1042/BCJ20200029.

Lee, J., Worrall, L.J., Vuckovic, M., Rosell, F.I., Gentile, F., Ton, A.T., Caveney, N.A., Ban, F., Cherkasov, A., Paetzel, M., and Strynadka, N.C.J. (2020). Crystallographic structure of wild-type SARS-CoV-2 main protease acyl-enzyme intermediate with physiological C-terminal autoprocessing site. Nat Commun 11, 5877. 10.1038/s41467-020-19662-4.

Lin, S., Chen, H., Ye, F., Chen, Z., Yang, F., Zheng, Y., Cao, Y., Qiao, J., Yang, S., and Lu, G. (2020). Crystal structure of SARS-CoV-2 nsp10/nsp16 2’-O-methylase and its implication on antiviral drug design. Signal Transduct Target Ther 5, 131. 10.1038/s41392-020-00241-4.

Littler, D.R., Gully, B.S., Colson, R.N., and Rossjohn, J. (2020). Crystal Structure of the SARS-CoV-2 Non-structural Protein 9, Nsp9. iScience 23, 101258. 10.1016/j.isci.2020.101258.

MacDonald, E.A., Frey, G., Namchuk, M.N., Harrison, S.C., Hinshaw, S.M., and Windsor, I.W. (2021). Recognition of Divergent Viral Substrates by the SARS-CoV-2 Main Protease. ACS Infect Dis 7, 2591–2595. 10.1021/acsinfecdis.1c00237.

Mengist, H.M., Fan, X., and Jin, T. (2020). Designing of improved drugs for COVID-19: Crystal structure of SARS-CoV-2 main protease M(pro). Signal Transduct Target Ther 5, 67. 10.1038/s41392-020-0178-y.

Narayanan, A., Narwal, M., Majowicz, S.A., Varricchio, C., Toner, S.A., Ballatore, C., Brancale, A., Murakami, K.S., and Jose, J. (2022). Identification of SARS-CoV-2 inhibitors targeting Mpro and PLpro using in-cell-protease assay. Commun Biol 5, 169. 10.1038/s42003-022-03090-9.

Passmore, L.A., and Russo, C.J. (2016). Specimen Preparation for High-Resolution Cryo-EM. Methods Enzymol 579, 51–86. 10.1016/bs.mie.2016.04.011.

Pettersen, E.F., Goddard, T.D., Huang, C.C., Couch, G.S., Greenblatt, D.M., Meng, E.C., and Ferrin, T.E. (2004). UCSF Chimera--a visualization system for exploratory research and analysis. J Comput Chem 25, 1605–1612. 10.1002/jcc.20084.

Punjani, A., Rubinstein, J.L., Fleet, D.J., and Brubaker, M.A. (2017). cryoSPARC: algorithms for rapid unsupervised cryo-EM structure determination. Nat Methods 14, 290–296. 10.1038/nmeth.4169.

Rut, W., Groborz, K., Zhang, L., Sun, X., Zmudzinski, M., Pawlik, B., Wang, X., Jochmans, D., Neyts, J., Mlynarski, W., et al. (2021). SARS-CoV-2 M(pro) inhibitors and activity-based probes for patient-sample imaging. Nat Chem Biol 17, 222–228. 10.1038/s41589-020-00689-z.

Rut, W., Lv, Z., Zmudzinski, M., Patchett, S., Nayak, D., Snipas, S.J., El Oualid, F., Huang, T.T., Bekes, M., Drag, M., and Olsen, S.K. (2020). Activity profiling and crystal structures of in-hibitor-bound SARS-CoV-2 papain-like protease: A framework for anti-COVID-19 drug design. Sci Adv 6. 10.1126/sciadv.abd4596.

Wang, Q., Wu, J., Wang, H., Gao, Y., Liu, Q., Mu, A., Ji, W., Yan, L., Zhu, Y., Zhu, C., et al. (2020). Structural Basis for RNA Replication by the SARS-CoV-2 Polymerase. Cell 182, 417–428 e413. 10.1016/j.cell.2020.05.034.

Xiao-chen Bai, G.M., Sjors H W Scheres (2014). How cryo-EM is revolutionizing structural biology. Trends in Biochemical Sciences Volume 40, Pages 49–57.

Zhang, L., Lin, D., Sun, X., Curth, U., Drosten, C., Sauerhering, L., Becker, S., Rox, K., and Hilgenfeld, R. (2020). Crystal structure of SARS-CoV-2 main protease provides a basis for design of improved alpha-ketoamide inhibitors. Science 368, 409–412. 10.1126/science.abb3405.

Zhao, Y., Zhu, Y., Liu, X., Jin, Z., Duan, Y., Zhang, Q., Wu, C., Feng, L., Du, X., Zhao, J., et al. (2022). Structural basis for replicase polyprotein cleavage and substrate specificity of main protease from SARS-CoV-2. Proc Natl Acad Sci U S A 119, e2117142119. 10.1073/pnas.2117142119.

Zhu, L., George, S., Schmidt, M.F., Al-Gharabli, S.I., Rademann, J., and Hilgenfeld, R. (2011). Peptide aldehyde inhibitors challenge the substrate specificity of the SARS-coronavirus main protease. Antiviral Res 92, 204–212. 10.1016/j.antiviral.2011.08.001.

